# Cholinergic calcium responses in cultured antennal lobe neurons of the migratory locust

**DOI:** 10.1101/2021.02.11.430736

**Authors:** Gregor A. Bergmann, Gerd Bicker

## Abstract

Locusts are advantageous organisms to elucidate mechanisms of olfactory coding at the systems level. Sensory input is provided by the olfactory receptor neurons of the antenna, which send their axons into the antennal lobe. So far, cellular properties of neurons isolated from the circuitry of the olfactory system, such as transmitter-induced calcium responses, have not been studied. Biochemical and immunocytochemical investigations have provided evidence for acetylcholine as classical transmitter of olfactory receptor neurons. Here, we characterize cell cultured projection and local interneurons of the antennal lobe by cytosolic calcium imaging to cholinergic stimulation. We bulk loaded the indicator dye Cal-520 AM in dissociated culture and recorded calcium transients after applying cholinergic agonists and antagonists. The majority of projection and local neurons respond with increases in calcium levels to activation of both nicotinic and muscarinic receptors. In local interneurons, we reveal interactions lasting over minutes between intracellular signaling pathways, mediated by muscarinic and nicotinic receptor stimulation. The present investigation is pioneer in showing that Cal-520 AM readily loads *Locusta migratoria* neurons, making it a valuable tool for future research in locust neurophysiology, neuropharmacology, and neurodevelopment.

## Introduction

Migratory locusts are of socio-economical importance as devastating agricultural pests, but they serve also in basic research as preparations for insect physiology, genomics, functional analysis of neural circuitry, and for understanding underlying mechanisms of development^1–8^. In the field of systems neurobiology, locusts have been instrumental to elucidate circuit mechanisms of olfactory coding in the insect brain^9^. The antennal lobe (AL) is the primary olfactory center in insects, resembling the olfactory bulb in mammals. Sensory input is provided by the olfactory receptor neurons (ORN) of the antenna, which send their axons into the AL^10^. In this primary olfactory center, they make synaptic connections with local interneurons and projection neurons. Similar to the architecture of the vertebrate olfactory bulb, the synaptic terminals of insect ORN interact with the dendrites of local interneurons and olfactory projection neurons in spherical neuropil compartments, termed glomeruli. Most investigated insects, like for example the genetic model Drosophila, employ a uniglomerular wiring strategy, in which ORN expressing the same olfactory receptor genes converge within the same glomerulus in the AL^11,12^. Locusts, with their roughly 1000^13^ glomeruli in the AL, provide a clear exception of this principle.

In this wiring strategy, each ORN axon branches in the AL, innervating multiple glomeruli, where they interface with both projection neurons (PN) and local interneurons (LN)^9,10,13,14^. Local interneurons, the most common of which being GABAergic and nitrergic^15^, also contact multiple glomeruli at once and play a modulatory role in this network^16,17^. The dendrites of each projection neuron, in turn, sample input from several glomeruli. The axons of the 830 olfactory projection neurons convey the output of the AL to the mushroom body and the lateral horn^10,13,16,18^. Since the olfactory system is very amenable both to anatomical and electrophysiological analysis^13,14,16–18^, extensive knowledge about odor encoding in the AL and the mushroom body has accumulated over the last four decades. Again, similar to the olfactory bulb, odor stimulation leads to oscillatory activity in the AL^17^. This synchronized electrical activity is generated through the recruitment of GABAergic inhibitory local neurons and reflected in the temporal dynamics of firing patterns of the projection neurons toward the mushroom body^17^.

Immunocytochemical investigations show that sensory neurons of the antenna express the acetylcholine synthetizing enzyme choline acetyltransferase, the vesicular acetylcholine transporter and a high affinity choline transporter^19–22^. Based on these investigations of several working groups, the majority of antennal ORN are thought to use acetylcholine as classical transmitter. There are two fundamentally different receptor types for acetylcholine, which are also present in insects. Ionotropic nicotinic acetylcholine receptors (nAChR) mediate mainly fast synaptic neurotransmission while metabotropic muscarinic acetylcholine receptors (mAChR) are thought to have modulatory roles via intracellular signal cascades, such as stimulation of phospholipase C, adenylate cyclase inhibition, and cation conductance changes^23–26^.

Unlike in the model organism Drosophila, where neural activity in defined of classes of neurons has been visualized using targeted expression of genetically encoded calcium indicators^27–29^, we are aware of only one calcium imaging study in the locust olfactory system. To estimate odor-evoked neuronal firing rate from Ca^2+^ signals, the calcium indicator Oregon-Green BAPTA-1 was injected into single projection neurons which were imaged using two-photon microscopy^30^. So far, cellular properties of neurons isolated from the circuitry of the olfactory system, such as transmitter-induced Ca^2+^ responses, have not been reported.

In this investigation, we aim to establish a procedure to characterize large numbers of olfactory system neurons by calcium imaging in dissociated cell culture. However, based on our own experience with cultured locust neurons of thoracic ganglia, there are difficulties in recording cytosolic signals with a number of commonly applied fluorescent calcium sensors in the acetoxymethyl ester (AM) form, due to rapid intracellular compartmentalization and dye extrusion. Here, we use the next generation calcium indicator Cal-520 AM to load AL neurons of *Locusta migratoria* and record their response to cholinergic stimulation. To distinguish between classes of local and projection neurons in culture, we label GABAergic and nitrergic LN^15,16^ and develop additional criteria from measuring soma size for appropriate identification. We find Ca^2+^ responses to cholinergic agonists in neurons of both classes. We also show crosstalk between intracellular signaling pathways mediated by nicotinic and muscarinic receptors. This opens new avenues to analyse transmitter-related properties of olfactory network components of a pest insect in a controlled environment. Moreover, it allows for a comparison of published cellular properties of AL neurons in the genetic model Drosophila^29^ to those of an insect relying on a different neuroanatomical wiring, but with well-investigated electrophysiology of the olfactory system^9^.

## Results

### Cell populations of the antennal lobe

To characterize neuronal phenotypes of the AL in the species *Locusta migratoria*, we started with a survey of its complete number of cells. We counted DAPI-labeled nuclei on frontal tissue sections, which were double-labeled for anti-acetylated tubulin (Fig.1a) and GABA (Fig.1b). Evaluation of the composite allowed for a comparison of nuclear size (Fig.1c) of the GABAergic LN (Fig.1d) with the remaining nuclei belonging to cells of undetermined transmitter phenotype. The AL comprised an overall mean of 1422 cells (neurons plus glia), including 83 GABAergic LN (Tab.1). These cell counts revealed no significant differences between males and females (Tab.1; p = 0.0823, p = 0.5368; Mann-Whitney test).

**Figure 1:**
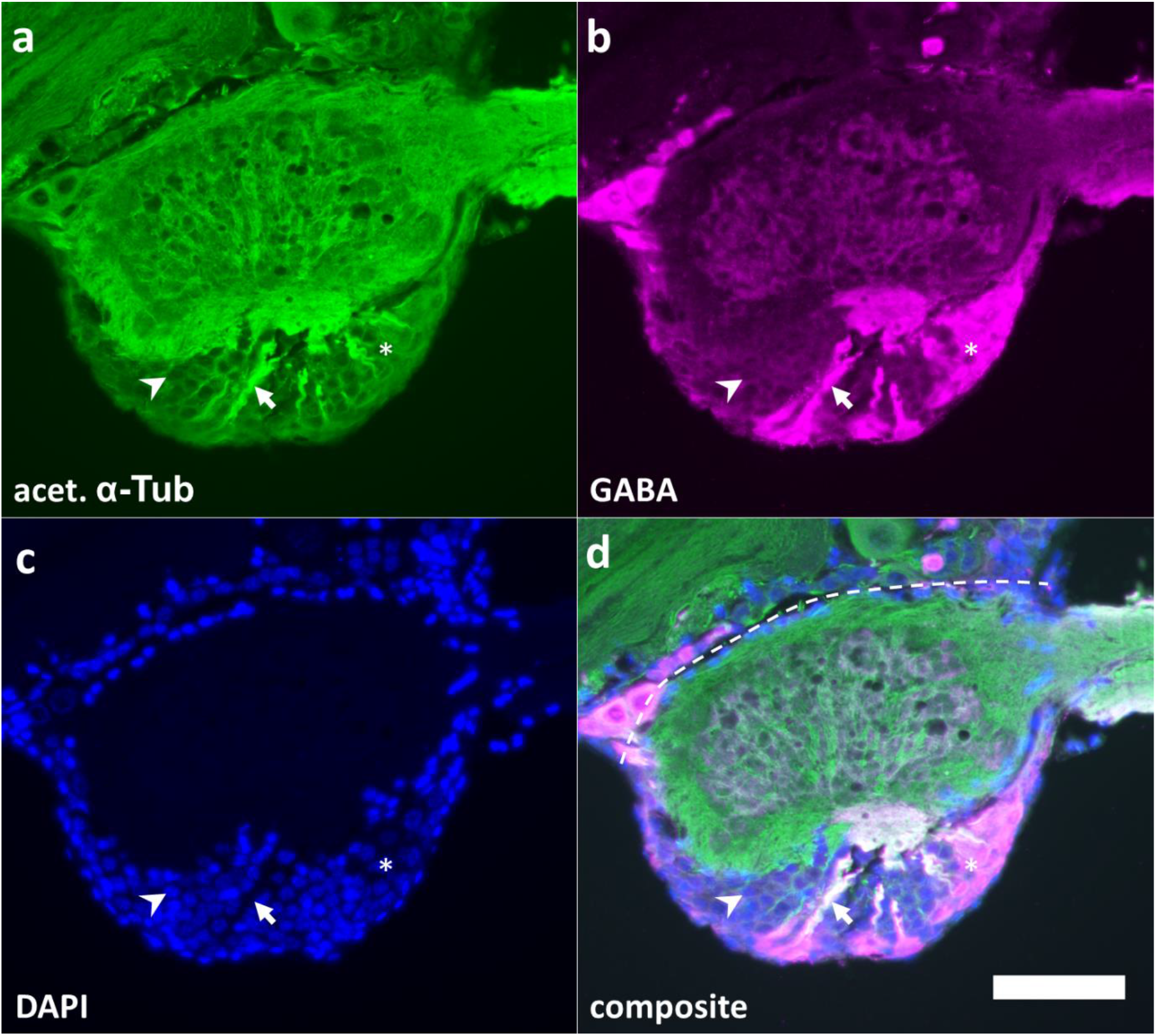
Frontal sections through the antennal lobe of *Locusta migratoria*. Sections were labeled using anti-acetylated α-tubuline (**a**) and anti-GABA (**b**) antisera, and the nuclear marker DAPI (**c**). GABA-IR local neurons (asterisk) were found in the frontal cell group of the antennal lobe. Somata of GABA-IR local neurons were typically larger compared to other cells (**c**). Most PN are also found in the frontal cell group, but had significantly smaller nuclei (arrowhead). In **d**, white color indicates colocalisation of GABA and acetylated α-tubuline. GABAergic LN and PN arborize into glomeruli (arrow). No arbors or connecting neurites were found on the dorsal side of the antennal lobe (segmented line). Scalebar = 100 μm.

**Table 1:**
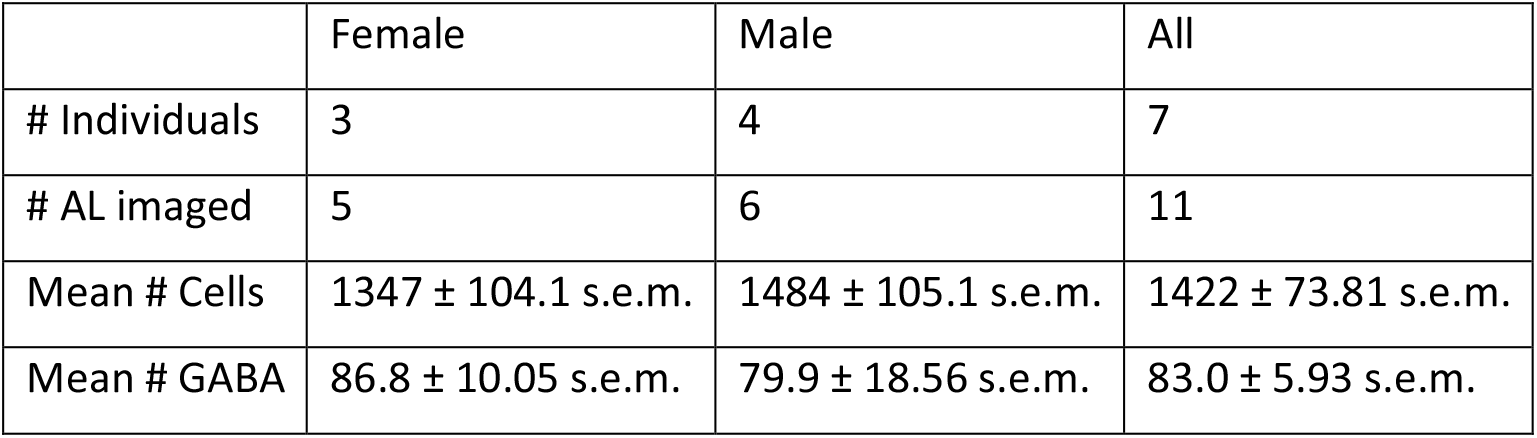
Comparison of nuclear countings from antennal lobe of female and male migratory locusts. Nuclei of 11 antennal lobes (AL) from seven individuals were analyzed. Nuclear counting comprises PN, LN, and glia. Mann-Whitney tests revealed, that nucleus counts of all cells and GABAergic cells was not significantly different between male and female locusts (p = 0.0823; p = 0.5368 respectively).

|  | Female | Male | All |
| --- | --- | --- | --- |
| # Individuals | 3 | 4 | 7 |
| # AL imaged | 5 | 6 | 11 |
| Mean # Cells | 1347 ± 104.1 s.e.m. | 1484 ± 105.1 s.e.m. | 1422 ± 73.81 s.e.m. |
| Mean # GABA | 86.8 ± 10.05 s.e.m. | 79.9 ± 18.56 s.e.m. | 83.0 ± 5.93 s.e.m. |

For physiological characterization, we chose to isolate and culture AL neurons in order to remove network interactions. In primary cell culture, we recovered 16.9 % (± 7.1 s.d. n = 29) of cells for immune- and cytochemical characterization of GABAergic and nitrergic phenotype^15^, indicative of local neurons. The majority of neurons, presumably PN, were unlabeled cells with small somata of about 15 μm in diameter (Fig.2 arrows). In line with a previous histological study^15^, GABA-IR (Fig.2a, b, c asterisks), NADPHd positive (Fig.2d, e, f asterisks), and double labelled cells LN were also present (Fig.2g, h, i asterisks). A fifth class comprised large neurons with soma diameters of about 20 μm and no determined transmitter phenotype, presumably belonging to LN (Fig.2k, l, m asterisks).

**Figure 2:**
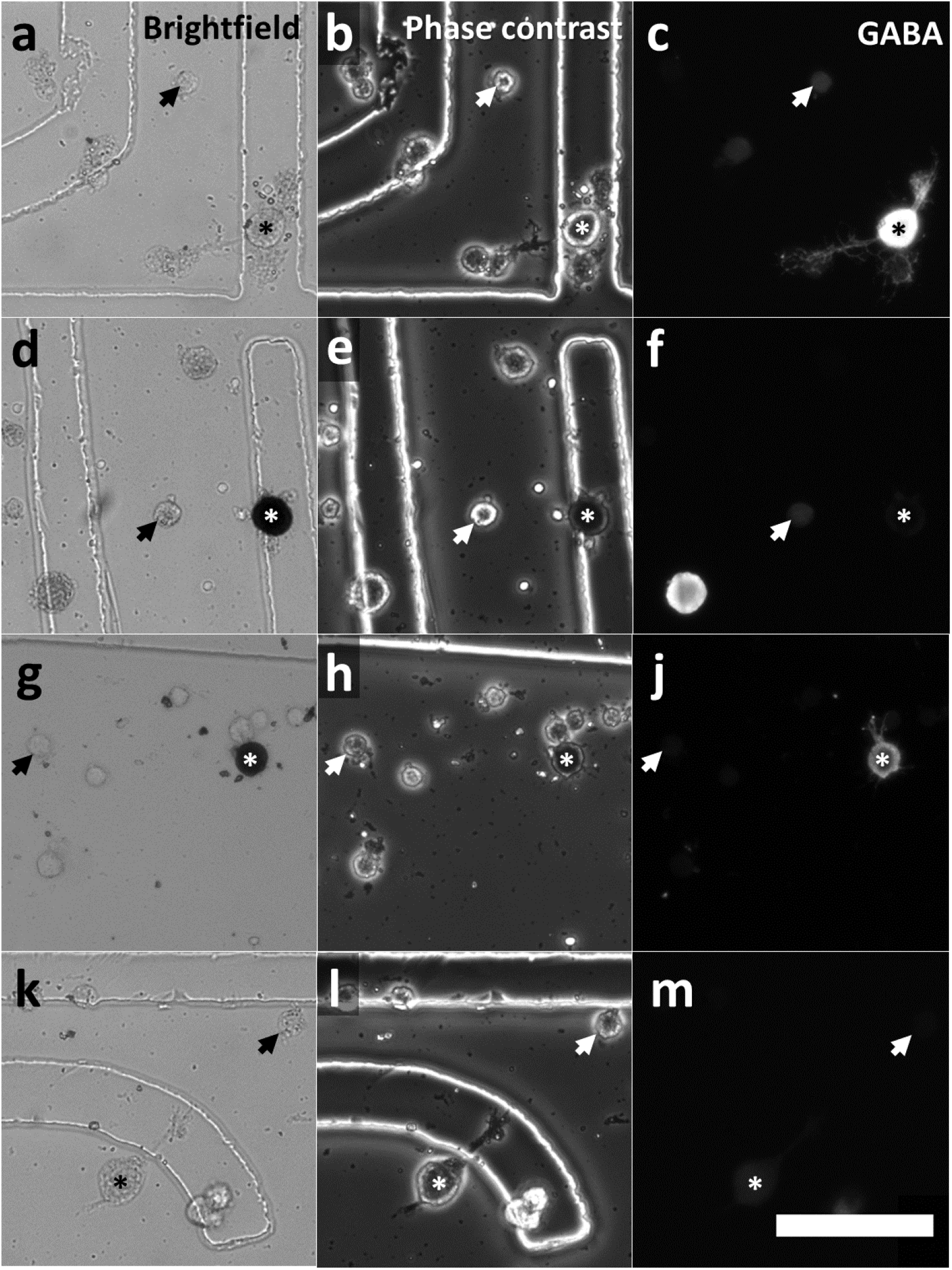
Cellular phenotypes of cultured antennal lobe neurons. Adherent neurons of different soma diameter were grown in petri dishes that contained a numbered location grid for proper cell identification after calcium imaging. Small neurons with soma diameter of about 15 μm are considered as PN (arrows), whereas large neurons of soma diameters above 19.61 μm are regarded to be LN (asterisk). NADPHd activity and GABA-IR was detected in large neurons. LN were classified into four types: exclusively GABAergic (**a-c**); NADPHd positive (**d**-**f**); GABA-IR and NADPHd positive (**g-j**); LN with no determined (ND) transmitter (**k-m**). Note that LN are generally larger in diameter, but not homogenous in size. Interneurons showing GABA-IR are common, but not morphologically distinct from other non GABA local neurons. LN expressing nitric oxide synthase were detected using NADPHd staining as marker(**d**, **g**). NADPHd staining tends to quench the fluorescence of GABA-IR local neurons (**j**). Large unlabeled LN showing neither NADPHd activity nor GABA-IR (**k-m** asterisk). Scalebar = 100μm.

To obtain a purposeful distinction between PN and LN, we initially quantified nucleus size from the paraffin sections and cell culture. To avoid any erroneous inclusion of other brain parts and perineural glia, we took care to evaluate only round nuclei in the anterior half of the AL^31^. Nuclei of GABA-IR neurons (Mean: 16.35 μm ± 0.11 s.e.m. n = 500) were significantly larger than nuclei of all sampled AL cells (Mean: 12.17 μm ± 0.07 s.e.m. n = 1716; p < 0.0001, Kruskal-Wallis test; Fig.3). Nuclear diameters of GABA-IR neurons showed a normal distribution, whereas those of all measured cells did not (Fig.3a & b; p = 0.66 and p < 0.0001 respectively; D’Agostino & Pearson normality test). Nuclear diameters of GABAergic LN appeared to be a normally distributed subgroup of all nuclei (Fig.3b i), since their mean diameter coincided with the plateau found in the tail of the distribution of all nuclei. For better illustration, we normalized relative frequencies of the GABAergic LN nuclear diameters to the total number of measured cells (Fig.3b).

**Figure 3:**
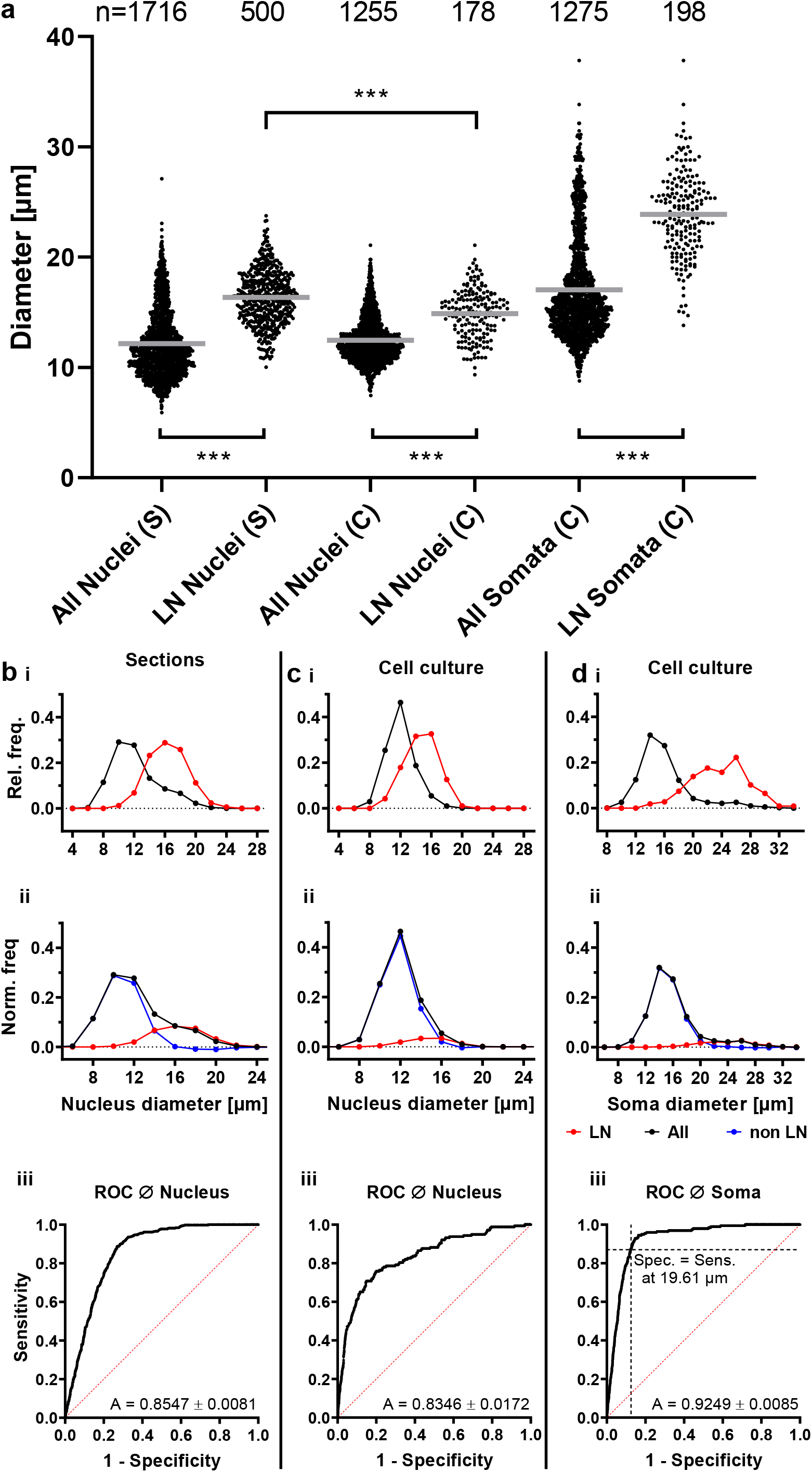
Discrimination of local and projection neurons based on nucleus and soma diameter. Individual size measurements of nuclei and somata in sections (S) and culture (C) including respective mean (gray line) are shown in **a**. Nuclear or soma diameters of labeled LN were significantly larger than respective pooled diameters of all neurons in sections and culture (p < 0.0001; Kruskal-Wallis test). Nuclear diameters of labeled LN were significantly smaller in culture than in sections (p = 0.0001, Kruskal-Wallis test), due to losses during dissociation and culturing. Relative frequencies of pooled nuclear diameters of all cells (including LN) in black, and labeled LN in red, as determined in sections (**b i)** and culture **(c i)**. **d i** shows relative frequencies of soma diameters of corresponding cultured neurons shown in **c**. In **b**-**d ii** frequencies were normalized to cell counts of all cells. Subtraction of normalized LN frequencies (red) from the pooled cell frequencies (black) resulted in a distribution of neurons without LN (non LN; blue). Non LN frequency at the mean nuclear/soma diameter was close to zero. This allowed estimation of the number of LN in sections (24 %) and culture (10.2 %). For binary LN/PN classification, ROC curves of the classifiers nuclear and soma diameter were plotted (**b**-**d iii**). The area under the ROC curve (A in **b**-**d iii**) reflects classifier suitability. For classification of cultured neurons, a threshold soma diameter for discrimination of LN from PN was derived at a value 19.61 μm, accounting for equal classifier specificity and sensitivity (i.e. equal false positive and false negative rate; **d iii**).

Assuming that all LN nuclei are of similar size, we analyzed whether nuclear diameters of GABA-IR cells could be used to estimate the percentage of LN (GABA-IR and other) in the AL. First, we calculated the normalized frequency of non LN nuclear diameter as the difference of the normalized frequencies of all cells and GABAergic LN. This uncovered the range of overlap in both distributions (Fig.3b ii). Both distributions overlap considerably at diameters between 10 μm and 12 μm. Intriguingly, the normalized frequency of the presumptive non LN nuclear diameters was close to zero at the mean LN nuclear diameter, which allowed us to calculate the percentage of LN in the AL. We counted all cells with nuclear diameters that were larger than the mean nuclear diameter of GABAergic LN (representing 50 % of all LN), doubled the outcome and divided by the total number of measurements. According to this calculation, roughly 24 % of AL cells are LN.

Based on nuclear diameter, PN and LN were not sufficiently distinguishable in culture. Due to losses of very large cells, nuclei of cultured LN (14.88 μm ± 0.17 s.e.m. n = 178) appeared to be significantly smaller than LN nuclei in sections (Fig.3a; p < 0.0001, Kruskal-Wallis test). Despite a statistically significant difference between nuclear diameters of all measured neurons and labeled LN (Fig.3a, p = 0.0001, Kruskal-Wallis test), the normalized frequency of non LN at mean LN nucleus diameter was not close to zero (Fig.3c ii). However, when soma diameter was used for LN classification, the normalized frequency of non LN at mean LN soma diameter was close to zero (Fig.3d ii). Another way to evaluate the suitability of a given classifier is the comparison of the area under respective receiver-operating characteristic (ROC) curves, in which a larger area indicates a better classifier. ROC analysis confirmed that soma diameter (Fig.3d iii; A = 0.924 ± 0.008 s.e.) is a better classifier than nuclear diameter (Fig.3c iii; A = 0.835 ± 0.017 s.e.) for estimation of the LN population. Therefore, soma diameters of labeled LN were used for the estimation of LN number in culture. Soma diameters of labeled LN were normally distributed, whereas diameters of all neurons were not (Fig.3a, d; p = 0.275 and p < 0.0001 respectively; D’Agostino & Pearson normality test). Diameters of all measured neurons (15.97 μm ± 0.10 s.e.m. n = 1275) were significantly different (p < 0.0001, Kruskal-Wallis test) to diameters of labeled LN (23.87 μm ± 0.87 s.e.m. n = 198). ROC analysis showed a classifier specificity of 0.94 at mean LN soma diameter. We calculated a percentage of 10.2 % LN in the cultures, indicating a preferential loss of LN over PN in the cell culture procedure. Additionally, the ROC analysis allowed to calculate a criterion for the distinction between PN and LN based on soma diameter. At a threshold diameter of 19.61 μm, the sensitivity was equal to the specificity on the ROC curve. Cells with soma diameters smaller or larger than this threshold were predicted as PN or LN respectively and, from now on, will be termed accordingly.

### Calcium imaging of cultured antennal lobe neurons

After unsuccessful attempts with other indicators (Suppl. Tab. S1), we reliably detected calcimycin-induced Ca^2+^ transients in cultured locust neurons by applying the calcium sensor Cal-520 AM. We selected regions on the dish that contained PN and LN using the previously established criterion. Following the imaging sessions, it was possible to verify a potential GABAergic or nitrergic phenotype of LN by immunofluorescence and cytochemistry.

Cultured cells responded with pronounced Ca^2+^ transients when stimulated with the unspecific cholinergic agonist carbachol (Fig.4a). Occasionally, they showed also spontaneous transients of lower magnitude (Fig.4a, Suppl. Fig. S2). Inhibition of cholinergic receptors with specific antagonists reduced the amplitude of responses, yielding a variety of outcomes (Fig.4).

**Figure 4:**
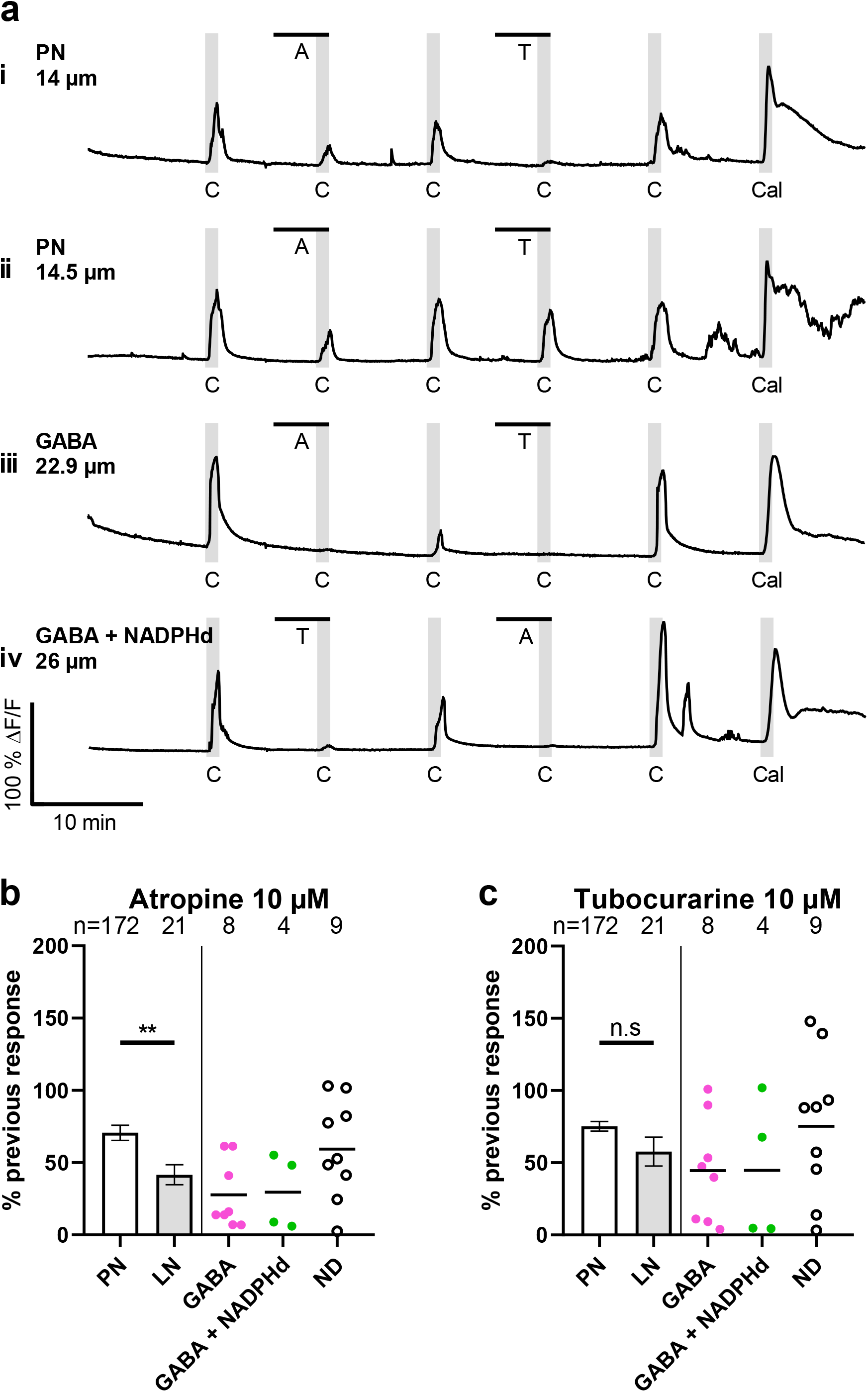
Reduction of carbachol-induced Ca^2+^ transients by co-incubation with cholinergic antagonists. At 10 μM, carbachol elicited Ca^2+^ transients, which were reduced when neurons were also exposed to 10 μM atropine or tubocurarine. Examples for Ca^2+^ signals of PN and LN are shown in **a i**-**iv**, illustrating the magnitude of reduced transients in individual neurons. The PN in **a i** showed stronger inhibition by the nicotinic antagonist tubocurarine than by the muscarinic antagonist atropine, whereas the PN in **a ii** showed a converse response. Labeled LN (GABA and GABA+NADPHd) showed generally stonger inhibition by antagonists up to a complete response abolishment (**a iii** & **iv**). This reduction of Ca^2+^ responses is summarized in **b** and **c** by normalization of amplitudes in the presence of antagonists to amplitudes prior to antagonist application. Bars show means ± s.e.m. of pooled PN and LN. Colored datapoints represent subtypes of individual LN. Atropine reduced responses of PN to 70.68 % (± 5.29 s.e.m. n=172), and to 41.64 % (± 6.95 s.e.m. n=21) for LN shown in **b**. These reductions were significantly different (p = 0.0048, Mann-Whitney test). As shown in **c,** tubocurarine reduced responses of PN to 75.25 % (± 3.39 s.e.m. n=172), and to 57.74 % (± 9.94 s.e.m. n=21) for LN, which were not significantly different (p = 0.0960, Mann-Whitney test).

In some PN, inhibition of muscarinic receptors with atropine resulted in a stronger reduction compared to nicotinic receptor inhibition with tubocurarine, but we also imaged PNs with converse responses (Fig.4a i and ii). Antagonists appeared to have stronger effects on labeled LN of GABAergic/nitrergic phenotype (Fig.4a iii and iv).

To account for the multitude of effects, we normalized Ca^2+^ amplitudes in the presence of inhibitors to the respective previous response to carbachol (Fig.4b). Atropine reduced carbachol responses in PN to 70.68 % (± 5.29 s.e.m. n = 172). We found twelve labeled LN showing GABA-IR, four of which were double-labeled for NADPHd activity. When pooled, atropine reduced LN responses to a mean of 41.64 % (± 6.95 s.e.m. n = 21) which was significantly different to PN response reduction (p = 0.0048, Mann-Whitney test). Tubocurarine reduced PN amplitudes to 75.25 % (± 3.39 s.e.m. n = 172), and the amplitudes of LN to 57.74 % (± 9.94 s.e.m. n = 21) of the previous response. We found no significant differences in response reduction by tubocurarine between PN and LN (p = 0.0968, Mann-Whitney test). These findings suggest that muscarinic and nicotinic acetylcholine receptor activation causes Ca^2+^ transients in AL neurons.

Next, we examined whether LN and PN show different responses to stimulation with nicotinic and muscarinic agonists (Fig.5, Suppl. Video S3). In range finding experiments, we determined that 1 μM nicotine and 10 μM pilocarpine elicit Ca^2+^ transients of similar magnitude in the majority of PNs (Fig.5a i). However, depending on the agonist and the individual neuron, responses were not entirely consistent in amplitude. Alternating nicotine and pilocarpine stimulations revealed three basic response types in PN with equal responses to both agonists, stronger responses to nicotine, and stronger responses to pilocarpine (Fig.5a i-iii). To illustrate whether these response types follow a certain distribution, we calculated the difference between the respective nicotine- and pilocarpine-elicited amplitudes (ΔN – ΔP), and plotted their relative frequency in a histogram (Fig.5b). Averaged over a large sample of 253 cultured neurons, the used concentrations elicited Ca^2+^ responses of similar peak amplitude in roughly one third of PN (Fig.5b-d). LN showed a different behavior towards the same experimental protocol.

**Figure 5:**
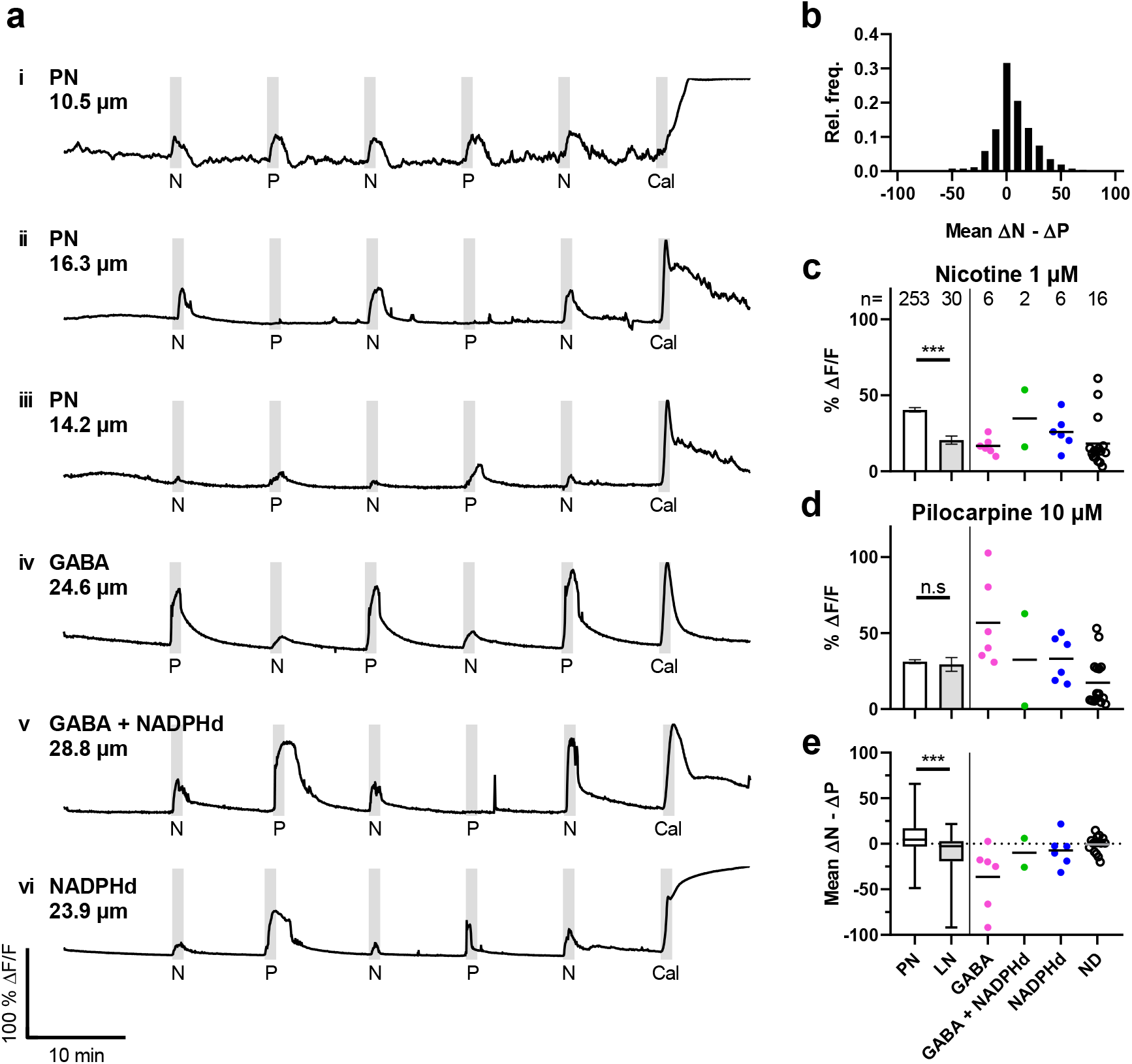
Nicotine and pilocarpine induced-Ca^2+^ transients of cultured antennal lobe neurons. Representative Ca^2+^ signals of PN (**i**-**iii**) and LN (**iv**-**vi**) are shown in **a**. PNs show three response types (equal **i**; stronger nicotinic **ii**; stronger muscarinic **iii**). A histogram (**b**) illustrates the distribution of these types as difference in response amplitude, with 31.6 % of PN showing equivalent responses. Mean amplitudes (± s.e.m.; normalized to calcimycin) of pooled PN and LN during stimulation with 1 μM nicotine (PN: 40.41 % ΔF/F ± 1.45 s.e.m.; LN: 20.51 % ΔF/F ± 2.69 s.e.m.) or 10 μM pilocarpine (PN: 31.24 % ΔF/F ± 1.27 s.e.m.; LN: 29.38 % ΔF/F ± 4.51 s.e.m.) are depicted in **c** and **d**. Colored data points represent subtypes of individual LN. During nicotinic stimulation, Ca^2+^ transients of PN were significantly larger than those of LN (p < 0.0001; Mann-Whitney test). When stimulated with pilocarpine no significant differences between PNs and LN were observed (p = 0.3825; Mann-Whitney test). Amplitude difference between nicotine and pilocarpine induced responses (ΔN - ΔP; **e**) is used to quantify the relationship between nicotinic and muscarinic responses in individual neurons (PN: 6.84 % ΔF/F ± 1.13 s.e.m.; LN: −9.77 % ΔF/F ± 4.17 s.e.m.). LN show a significant difference to PNs (p < 0.0001; Mann-Whitney test).

Labeled LN frequently showed larger Ca^2+^ transients during muscarinic than nicotinic stimulation (Fig.5a iv-vi). However, nicotine and pilocarpine responses were not consistent in amplitude, similar to responses in PN. On average, nicotinic responses of LN subtypes were smaller in amplitude than nicotinic responses of PN (Fig.5c). Pooling of these LN subtypes resulted in a mean nicotinic amplitude of 20.51 %ΔF/F (± 2.69 s.e.m. n = 30), which was significantly different to the respective response amplitude of PN (Mean: 40.41 %ΔF/F ± 1.45 s.e.m. n = 253; p < 0.0001, Mann-Whitney test). Pilocarpine responses of NADPHd positive LN were on average similar in amplitude to PN (Fig.5d). In contrast, pilocarpine responses of exclusively GABAergic LN (56.7 %ΔF/F ± 11.70 s.e.m. n = 6) were larger on average than PN amplitudes, whereas amplitudes of ND local neurons were lower (17.36 %ΔF/F ± 3.94 s.e.m. n = 16). This difference resulted in a mean response amplitude of 29.38 %ΔF/F (± 30.30 s.e.m. n = 2) when LN were pooled, which was not significantly different to PN amplitude (p = 0.3825, Mann-Whitney test).

To quantify the relationship between nicotinic and muscarinic responses in individual neurons, we evaluated the difference of corresponding response amplitudes (ΔN - ΔP) (Fig.5e). LN (Mean: −9.77 %ΔF/F ± 4.17 s.e.m. n = 30) were significantly different (p < 0.0001; Mann-Whitney test) from PN (Mean: 6.84 %ΔF/F ± 1.13 s.e.m. n = 253). This indicated that LN show stronger responses to pilocarpine than to nicotine. Furthermore, exclusively GABAergic neurons (Mean: −36.33 %ΔF/F ± 14.42 s.e.m. n = 6) seem to form a subset of local neurons that are characterized by larger Ca^2+^ transients in response to pilocarpine.

Cholinergic neurotransmission can be modulated by crosstalk of intracellular signaling pathways triggered by muscarinic and nicotinic receptor activation^32^. To detect potential interactions, we utilized a shortened protocol with stimulations every 2 min for 15 s. The protocol consisted of two 1 μM nicotine pulses, one pulse of a co-applied nicotine/pilocarpine mixture (1 μM and 10 μM respectively), another test pulse of nicotine, and a pulse of 10 μM pilocarpine. As a control, we replaced the nicotine/pilocarpine pulse with a nicotine pulse in half of the experiments to determine whether nicotinic stimulations yielded consistent responses. Projection neurons showed similar transients during each nicotinic stimulation, both in co-application as well as in the control (Fig.6a i-iii, b). On average, we found no significant differences between amplitudes of the control and the respective amplitudes of the co-application protocol (Fig.6b; lowest p = 0.6986 N2 vs. N2; Kruskal-Wallis test). In the example of Fig.6a i, co-stimulation elicited in the PN a larger transient than nicotine alone, whereas pilocarpine alone induced a transient of similar magnitude. This was not the case for all PN (Fig.6a ii-iii). To check whether such increased responses can be explained by a simple addition of pilocarpine and nicotine-induced Ca^2+^ transients during co-application, we calculated the sum of the nicotinic response (N2) prior to co-application and the respective muscarinic response (P) for each neuron. We found no significant alterations in nicotinic responsiveness after co-stimulation in PNs (p > 0.9999; Friedman test). Friedman’s test for paired data revealed a significant difference between the sum of nicotine and pilocarpine responses and the respective co-application (p < 0.0001).

**Figure 6:**
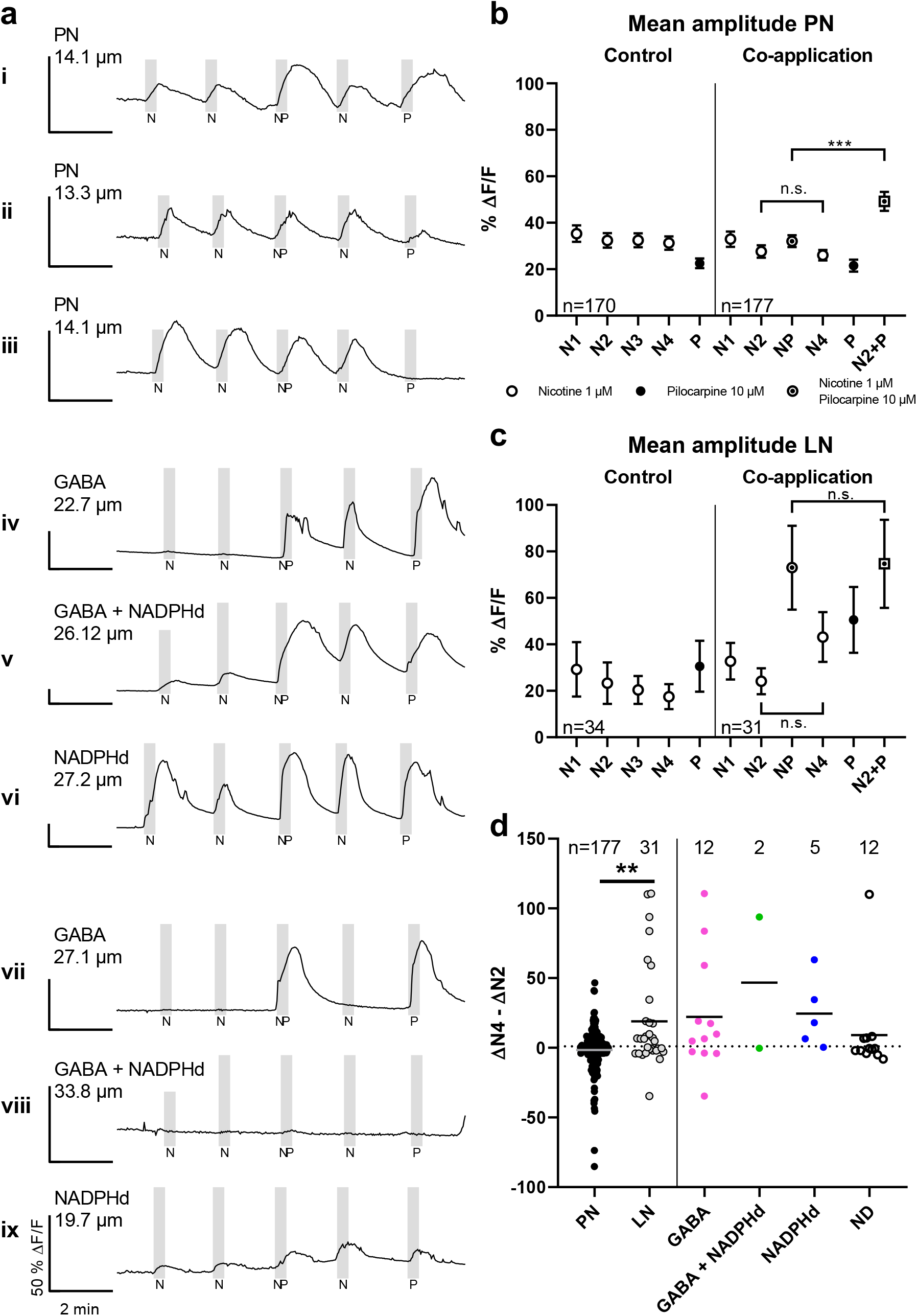
Ca^2+^ response alterations by interaction of muscarinic and nicotinic stimulation. To test for lasting response alterations, a series of two 15s pulses of 1 μM nicotine, one pulse of a 1 μM nicotine and 10 μM pilocarpine mix (no pilocarpine in controls), one nicotine test pulse and one pilocarpine pulse were given in consecutive order. **a** shows examples of Ca^2+^ signals in the co-application protocol. Note that scale varies for better visibility, but all represent 50 % ΔF/F. **a** shows various traces of PN (**i**-**iii**) and different LN types (**iv**-**ix**). Nicotinic responses were enhanced after co-application in several LN of all types (**a iv**-**vi**), but not in PN. This response enhancement was not found in all of the sampled LN (**a vii**-**ix**). **b** and **c** depict mean Ca^2+^ responses (± s.e.m.) of PN and LN normalized to calcimycin response. Responses to nicotine (empty circle), pilocarpine (filled circle), or nicotine/pilocarpine mix (circled dot) are shown in order of application. The predicted N2+P value (squared dot) was calculated as the sum of individual nicotine and pilocarpine responses during co-application. These predicted responses were significantly different to the respective co-application response in PNs (p < 0.0001; Friedman test), but not in LN (p = 0.1802; Friedman test). In PNs as well as LN (**b** & **c**), nicotinic amplitudes before and after co-application were not significantly different to each other (p > 0.9999 respectively; Friedman test). To account for variability in baseline responsiveness, amplitude differences of nicotinic stimulations after (N4) and prior to co-application (N2) of individual PN (black), LN (gray) and their respective subtypes were calculated (**d** line represents mean). Amplitude differences of LN were significantly different to those of PNs (p = 0.0073; Mann-Whitney test), demonstrating an enhancement of nicotinic responses after co-application of the muscarinic and nicotinic agonists.

Local neurons, however, responded notably different than PN. Labeled LN exhibited rather varied responses. Whereas some showed enhanced nicotinic responses after co-application compared to prior nicotinic responses (Fig.6a iv-vi), others showed no or weak responses to nicotine throughout the experiment (Fig.6a vii-ix). We found enhanced nicotine responses after co-application in all three labeled LN subtypes. These effects were also noticeable in larger mean response amplitude of the 31 pooled LN (Fig.6c). A significant difference (p = 0.0053; Kruskal-Wallis test) was only apparent between the third nicotine stimulation in the control (N3) and the nicotine/pilocarpine stimulation in the co-application (NP). The experimentally determined signal amplitude of co-application corresponded nicely to the predicted calculated sum, as no significant difference between co-application and the sum of nicotine and pilocarpine responses was found (Fig.6c; p = 0.1802; Friedman test). This indicated that for LN, the response amplitude during co-stimulation is indeed the result of nicotinic and muscarinic Ca^2+^ transient addition.

When we examined whether a co-stimulation of nicotinic and muscarinic receptors has notable effects on following nicotinic stimulations for the pooled LN (Fig.6c), we measured a slight increase in mean amplitude without statistical significance (p > 0.9999; Friedman test). This is likely accounted for by heterogeneity in response levels and in contrast to findings in individual traces. To provide a more detailed analysis of the enhanced response effect in LN after co-stimulation, and to compare it to responses in PN, we calculated the difference of nicotinic amplitudes after and prior to co-application (ΔN4 – ΔN2) in order to normalize for baseline responsiveness to either agonist (Fig.6d). This resulted in tailed distributions of amplitude differences, reflecting response variability in individual PN and LN. Most PN showed no or little change in amplitude (Mean: −1.56 %ΔF/F ± 1.17 s.e.m. n = 177). The distribution of PN amplitude differences was tailed in both directions (Fig.6d). In contrast to PN, the distribution of LN had a larger tail towards stronger nicotine responses after co-application (Mean: 19.02 %ΔF/F (± 6.58 s.e.m. n = 31). Amplitude differences of LN and PN were significantly different to each other (p = 0.0073; Mann-Whitney test), revealing an enhanced responsiveness of LN to nicotinic stimulation after co-application of a muscarinic agonist.

## Discussion

In this investigation, we characterized AL neurons of *Locusta migratoria* neurons by calcium imaging in cell culture, focusing on the classical transmitter of the ORN, acetylcholine (ACh). We were motivated by other investigations that used calcium indicators in the AM form for recording ACh-induced cell responses in primary cultured insect neurons^33,34^, and succeeded to obtain Ca^2+^ transients in the locust by loading the indicator Cal-520 AM (Fig.4; Suppl. Tab. S1). We showed that both PN and LN respond to nicotinic stimulation (Figs.4; 5). The majority of both types of neurons were also responsive to muscarinic stimulation, with local neurons as the more sensitive cellular phenotype (Fig.5). Simultaneous nicotinic and muscarinic stimulation of subsets of local neurons caused an enhanced Ca^2+^ response to a subsequent nicotine stimulus, indicating an intracellular interaction triggered by the activation of the two classes of cholinergic receptor types (Fig.6).

A prominent subset of the LN in the locust AL use GABA as classical neurotransmitter^9,15,16,35^. We utilized this information to distinguish between PN and LN in cell culture (Fig.3), without preceding labeling. Using ROC curves, we could approach the binary classification problem in culture by analyzing the prediction of the transmitter phenotype GABA from the soma diameter (Fig.3 iii). We settled for a criterion on the ROC curve, where the sensitivity is equal to the specificity, considering somata larger than a diameter of approximately 20 μm as LN and the PN as smaller. This procedural method, although afflicted with a small risk of inaccuracy, made it unnecessary to label neurons with tracers in the experimental animals prior to dissociation. Nonetheless, to confirm the GABAergic phenotype, all calcium imaging sessions were followed by labeling for GABA-IR.

Based on the simple assumption of a similar nuclear size distribution of GABA-positive LN and other unlabeled LN, we could also estimate the total number of LN from size measurements in tissue sections through the AL. These calculations excluded the glial cells, which make up 30% of locust brain cells^31^. Using the size distribution of DAPI-labeled nuclei of GABAergic neurons as proxy for all LN (Fig.3 a,b) we estimate that 24 % of the AL neurons are LN. This corresponds nicely to the 26% LNs which were reported for Schistocerca^9^. While the AL of Schistocerca seems to contain around 1130 neurons in total^9,16^, we estimated about 1000 neurons in the AL of *Locusta migratoria*. The number and the ratio between LN and relaying PN is considered to be an important factor in the oscillatory transient synchronization of AL output during odor encoding in the intact animal^9^. Pharmacological blocking of the inhibitory synapse between the GABAergic LN to the PN abolishes spike synchronization without affecting the response pattern of each individual neuron^35^.

For evaluation of responses to the cholinergic stimulation in cell culture, we chose the peak amplitude of the Ca^2+^ transients. In the diagrams we present separately the data for LN, that were immuno- and cytochemically (GABA, NADPH, double-labeled) characterized and the LN with not determined transmitter phenotype (ND). Due to a smaller sample size of neurons with confirmed neurotransmitter phenotype, we chose to perform statistical tests only between all PN and LN, as defined by the ROC method (Figs.4–6).

Similar to vertebrates, there are two fundamentally different receptor types for ACh in insects. Ionotropic nicotinic AChR for fast synaptic neurotransmission and metabotropic muscarinic AChR with modulatory functions via intracellular signal cascades^23–26^. In Locusta, five cDNA clones that encode nAChR subunits have been identified^36^. As for all insect nAChR subunits, functional expression in heterologous systems is not easy to achieve, but has been successful for an alpha-like subunit of Schistocerca in Xenopus oocytes^37,38^ and a novel beta subunit^39^. Modulation of ACh release from synaptosome preparations provided biochemical evidence for muscarinic receptors in the CNS of locusts^40,41^. The gain of the monosynaptic connection between the cholinergic locust wing stretch receptor and an identified wing depressor motoneuron is modulated by muscarinic autoreceptors on the presynaptic terminals of the stretch receptor itself and/or on muscarinic receptors of GABAergic interneurons^42,43^. In intact Drosophila, muscarinic stimulation and blocking of mAChR diminish or enhance Ca^2+^ transients of PN respectively during odor presentation^29^. In the same study, mAChR have been demonstrated to increase LN excitability as a means to counteract short-term depression at the ORN to LN synapse. Here we show that, compared to PN, especially GABAergic LN generate considerably stronger Ca^2+^ transients in response to muscarinic stimulation, whereas their respective nicotinic response appears to be lower than that of PN. However, unlike in Drosophila where PN do not express mAChR^29^, muscarinic stimulation elicited clear-cut Ca^2+^ transients in most locust PN (Fig.5). Even though both the spiking PN and non-spiking LN receive cholinergic input from ORN axon terminals in the glomeruli, both neuron types may react differently given their role in the olfactory circuit. This was confirmed by comparing Ca^2+^ transients of LN and PN in response to nicotinic and muscarinic agonists and pharmacological inhibition of their corresponding receptor types.

Surprisingly, co-stimulation enhanced subsequent nicotinic responses in some LN that showed no or rather small Ca^2+^ signals to nicotine prior to co-stimulation (Fig6a iv, v). The mechanistic underpinnings of these cellular phenomena and whether they are also involved in an increase in electrical excitability remain to elucidated. After stimulation with nicotine, the intracellular Ca^2+^ concentration can increase via influx through voltage-gated Ca^2+^ channels, through opening nAChR, and potentially by calcium-induced Ca^2+^ release from intracellular stores. Combined voltage-clamping and calcium imaging of dissociated somata from Locusta thoracic ganglia show that the influx through nAChR contributes to about 25 % of the Ca^2+^ signal after carbachol stimulation^44^. Calcium ion fluxes do not only change rapidly the membrane potential, but can also regulate signal transduction pathways on a longer timescale. Intracellular Ca^2+^ accumulation may thus cause lasting plastic changes in the cell biology of neurons. For example, Campusano et al.^45^ decribed a form of neuronal plasticity for cultured Drosophila Kenyon cells, in which a brief conditioning pulse of nicotine causes a decrease in responsivenes to a second nicotine application 4 h later. In contrast, we detected an enhancement in nicotine-induced Ca^2+^ responses after co-stimulation with nicotine and pilocarpine that seems to be a distinctive feature of a group of local neurons including some of the GABAergic LN (Fig.6d). The cellular memory of this form of plasticity lasts for at least 2 minutes. Potentially, neuromodulation by muscarinic receptors may upregulate the fast synaptic transmission via nicotinic receptors in the intact AL^29^.

A subset of LN releases NO in a calcium-dependent manner that stimulates the formation of cGMP in the same or other LN^15,46,47^. In our calcium imaging study, we also found examples of these LN which were responsive both to stimulation with nicotine and pilocarpine (Fig.5a v vi, c-e). Further investigations must show how increases in cGMP levels may in turn regulate intracellular Ca^2+^ levels and neuronal excitability.

We have also found spontaneous Ca^2+^ transients mainly in PNs that occured prior to carbachol stimulation, but also later at regular intervals between stimulation events. Since the shown traces in the Suppl. Fig. S2 were not from neurons linked by neurites, these transient do not reflect network activity. Rather they reflect cell intrinsic oscillations of similar frequency and amplitude (Suppl. Fig S2 a) and also irregular fluctions (b) of Ca^2+^ concentrations. In Drosophila cell cultures, spontaneous Ca^2+^ transients are also generated by Kenyon cells and about 60% of all central brain neurons^48^. Whether neuronal Ca^2+^ transients are present in the developing and adult locust nervous system remains a challenging question for future investigations.

So far, calcium imaging of locust neurons required intracellular injection of calcium indicators^30,44,49^, making simultaneous calcium imaging of large cell populations difficult. However, Isaacson and Hedwig described an innovative technique, in which calcium indicators were introduced via iontophoresis through the nerve sheath to multiple cells^50^. This method appears technically feasible for easily accessible nerves, but may be rather difficult to apply for loading neurons of tracts running in the depth of the brain, such as the antennal lobe tract. The present investigation is pioneer in showing that Cal-520 AM readily loads *Locusta migratoria* neurons, making it a valuable tool for future research in locust physiology. The genome of the migratory locust had been sequenced a few years ago^1^ which, alongside advancements in genome editing like CRISPR-Cas9, rekindled the interest in research regarding the locust olfactory system^51^. Moreover, recent investigations introduced locusts as helpful organisms for research on axonal regeneration in the olfactory system^52^ and as predictive test system for developmental neurotoxicity in humans^53,54^. Both lines of research will require information how locust neurons exposed to axonal injury or developmental-toxicants respond to these challenges in terms of Ca^2+^ concentration changes. Taken together, the possibility of calcium imaging of selected neuronal populations in a defined environment has the potential to unravel new insight into the cellular properties of the olfactory system of a devastating pest insect, which is also a fruitful preparation for basic neurobiology.

## Methods

Locusts (*Locusta migratoria*) from our crowded culture were used for cell culture of antennal lobe neurons and paraffin sectioning. Locust brains were dissected in sterile Leibovitz’s L15 containing 1 % penicillin/streptomycine (Invitrogen). All chemicals were purchased from Sigma, unless noted otherwise.

### Primary antennal lobe neuron culture

For primary antennal lobe culture, the cell culture protocol of Kirchhof and Bicker^55^ was modified. A minimum of ten brains of an equal number of male and female fifth instar larvae were dissected at room temperature. For optimal culturing of AL neurons, we used brain tissue from animals 1 - 3 days after molting. Brains were desheathed, partially lifting the AL away from the protocerebrum, allowing for easy removal with sharp forceps (see Suppl. Fig. S4).

In paraffin sections, we found no neuronal connections between the dorsal side of the antennal lobe and the protocerebrum (Fig.1 d, segmented line). Therefore, it is very unlikely that the removal of the antennal lobes removes a significant number of non antennal lobe neurons.

AL were incubated for 45 minutes in in sterile collagenase/dispase (1mg/ml in PBS, Roche), which was subsequently removed by five 5 minute washing steps with L15 medium. Tissue was mechanically dissociated by gentle up and down pipetting using a 200 μl siliconized pipette tip (VWR cat No. 53503-794). The resulting suspension was seeded in small 20 - 50 μl droplets on ibiTreat μ-Dishes (Ibidi cat No. 81156). Cells were allowed to adhere for 40 min, cultured at 30 °C in 3 ml L15 medium, and washed thoroughly every two days. To select for neurons, cells were kept in culture for at least seven days, since glia is not as viable in culture and dies within one week^31,55^.

### Calcium imaging

After unsuccessful attempts with several other calcium indicators (see Suppl. Tab. S1), we detected calcimycin elicited Ca^2+^ transients in cultured locust neurons by applying the calcium sensor Cal-520 AM. Prior to loading, cultured neurons were washed for 10 min with L15 medium (supplemented with 10 mM HEPES, adjusted to pH 7.3; Roth). Optimal loading of AL neurons was achieved with 8 μM Cal-520 AM calcium indicator (stock solution 8 mM in DMSO, Abcam cat No. ab171868) in L15/HEPES and an incubation time of 90 minutes at 30 °C. For laminar superfusion of chemicals, a Warner RC-37W perfusion insert was fitted into the culture dish. This setup was mounted on a Zeiss Axiovert 200 inverse epifluorescence microscope and connected to an open peristaltic perfusion system, delivering a constant stream of L15/HEPES at 4 ml/min.

The indicator fluorescence was detected at excitation with 475 nm light (Colibri 7; filter set 44; ZEISS). To reduce photobleaching and phototoxicity, images were taken every two seconds at 2 % light source intensity, 4×4 binning (Axiocam 503 mono; Zeiss), 2x signal amplification, and an exposure time of 500 ms. For image acquisition ZEN 2.6 Imaging software (ZEISS) was used.

All cholinergic agonists and antagonists were dissolved and diluted in L15/HEPES. Carbachol, nicotine, pilocarpine, and the calcium ionophore calcimycin (5 μM, stock solution 5 mM in DMSO) were applied via a 3-way valve which was installed between the medium reservoir and the peristaltic perfusion pump (mdx-biotechnik cat No. TL/150). Atropine and tubocurarine were applied by switching from the main L15/HEPES reservoir to a separate reservoir containing the antagonist solutions. For antagonist experiments (Fig.4), we superfused cultured antennal lobe neurons every 10 min for 1 min with 10 μM of the nonspecific cholinergic agonist carbachol. Using carbachol instead of acetylcholine avoided rapid breakdown by catalytic activity of acetylcholinesterase which is expressed on the soma surface of the majority of cultured locust neurons^56^. To probe for nicotinic or muscarinic components in the Ca^2+^ transients, every other carbarchol stimulus was applied in the presence of equimolar concentrations of an antagonist. We chose the muscarinic antagonist atropine and the nicotinic antagonist tubocurarine, which were pre-incubated for 5 min prior to carbachol stimulation. For agonist experiments (Fig.5), we stimulated cultured neurons with alternating 1 min nicotine and pilocarpine superfusions separated by 9 min washout periods. For the experiments shown in Fig.6, a shortened protocol with 15 second stimulations with the appropriate agonists and 105 seconds of washout was used. In co-application experiments, following agonists were applied in consecutive order: 1 μM nicotine; 1 μM nicotine; 1 μM nicotine + 10 μM pilocarpine; 1 μM nicotine; 10 μM pilocarpine. For the respective control, the nicotine pilocarpine mix was replaced by 1 μM nicotine.

### NADPH-diaphorase staining and GABA-immunocytochemistry of cell culture

For identification of the GABAergic and nitrergic phenotype of LN, cells were processed for GABA-IR and NADPH-diaphorase (NADPHd) activity^46^. After image acquisition, cells were fixed in cold GPFA (4 % paraformaldehyde dissolved in PBS with 0.1 % glutaraldehyde) on ice for 20 min, and rinsed with PBS (10 mM sodium phosphate, 150 mM NaCl, pH 7.4; Roth). Subsequently, fixed cells were washed two times in PBS-T (0.1 % Triton X-100; Roth) and once in 50 mM Tris HCl (pH 7.8, 0.1 % Triton; Roth). NADPHd activity was visualized by incubating with 50 mM Tris HCl containing 0.24 mM β-NADPH and 0.24 mM nitroblue tetrazolium for 20 min at room temperature, until stained LN acquired a dark violet color. Afterwards, cells were permeabilized with 0.3 % saponine in PBS-T for 20 min, washed and blocked with 5 % normal goat serum (Linaris) in PBS-T. The primary rabbit antibody against GABA (1:5000 in blocking solution; Sigma cat No. A2052) was incubated overnight at 4 °C. After three washing steps in PBS-T, cells were incubated with secondary biotinylated goat anti-rabbit (1:333 in blocking solution; Vector cat No. BA-1000) for at least 4 h at room temperature, followed by three more washing steps. GABA-IR was visualized by incubation with streptavidin conjugated Cy3 (1:333 in PBS-T) for at least 4 h at room temperature. Nuclei were labeled with 0.05 μg/ml 4’,6-Diamidino-2-Phenylindole (DAPI) simultaneously with the GABA-IR detection. For estimation of cell populations within the antennal lobe culture, we measured soma and nuclear diameters of cells within the field of view, as well as additional labeled LN outside the imaging frame. Since neurons can assume ellipsoid or spindle like shapes, we measured the minimal and the maximal diameter of somata and nuclei for each cell and used the mean.

### Paraffin sections and GABA-immunofluorescence

Paraffin sectioning of locust brains (Tab.1) was performed as described by Stern et. al.^57^. Briefly, brains were dissected in L15 medium, fixed in GPFA for 30 min at room temperature, rinsed in PBS, dehydrated, embedded in paraffin (Roth), sectioned at 14 μm, and mounted on chromealum/gelatin coated slides. Sections were rehydrated and washed in PBS. Immunolabeling was performed as in culture, including an additional labeling against acetylated-α-tubuline (monoclonal, 1:1000 in blocking solution; Sigma cat No. T6793), which was detected with Alexa488 conjugated goat anti-mouse (1:250 in blocking solution; Invitrogen cat No. A11001). A GABA-IR detection system (secondary antibody and streptavidin-Cy3) was used at 1:250 respectively, and DAPI was used at 0.1 μg/ml. Slices were washed, and cleared in glycerol. Nuclei within the antennal lobe were counted and evaluated according to the Abercrombie correction^58^. For determination of nuclear diameters, 1716 randomly selected cells were measured. Since nuclei of GABA-IR cells in the antennal lobe were generally larger in diameter, GABA-IR cells were counted and cell number was corrected separately, using diameters of 500 GABA positive cells. These measurements served to estimate the population of LN within the intact antennal lobe.

### Analysis of AL neuron types

To analyze cell populations, we generated histograms of nuclear and soma diameters (Fig.3 b-d i). Histograms shown in Fig.3 b-d ii were normalized to the nucleus/soma diameter distribution of all measured cells, which include LN as well as PN. The diameter distribution of non LN was calculated by subtracting the normalized frequencies of GABA-IR neurons from the normalized frequencies of all measured cells (Fig.3 b-d ii). Since the normalized frequency of non LN at the mean LN diameter is close to zero, LN percentage was calculated as follows. All cells with nucleus/soma diameters that were larger than the mean LN nucleus/soma diameter, representing 50 % of the total LN population, were counted, doubled and divided by the total number of cells. ROC curves were generated using the Wilson/Brown method in GraphPad Prism 7, based on nucleus/soma diameters obtained from sections and from cultured cells.

### Analysis of calcium imaging data

Fluorescence of individual Cal-520 AM loaded cells was measured in Fiji^59^ using manually fitted ROIs for each cell. Background fluorescence at each given frame was subtracted from individual cellular fluorescence. Baseline pre-treatment fluorescence (F) was measured 10 min prior to the first drug application. The change of fluorescence relative to baseline (ΔF/F) was calculated and normalized to the peak amplitude of the signal from stimulation with the calcium ionophore calcimycin (% ΔF/F). The amplitudes of Ca^2+^ responses to pharmacological stimulation (e.g. ΔN for nicotinic stimulation) were calculated as the difference of peak % ΔF/F and baseline fluorescence for each cell within a stimulation period. Cells with changes in fluorescence above 110 % relative to calcimycin were considered as damaged and thus excluded form data analysis. We did not apply this criterion for experiments of Fig.6 to preserve increased transients beyond the calcimycin maximum. In this experiment, we calculated a predicted value for co-application by simply adding amplitudes of the second nicotinic stimulation (N2) to the respective pilocarpine amplitude in each cell (Fig.6 b). This was necessary, since the first nicotinic stimulation (N1) tended to elicit slightly larger Ca^2+^ transients in control as well as in co-application experiments, and the nicotinic stimulation after co-application (N4) could be affected by the previous stimulation of mAChRs.

### Statistical analysis

Non-parametric tests were used for statistical analysis, since not all datasets showed normal distribution. All tested comparisons were considered significant at alpha < 0.05. Kurskal-Wallis tests for comparison of multiple datasets were performed with Dunn’s correction. Two tailed Mann-Whitney tests were used for comparing datasets with two groups (i.e. male vs. female or LN vs. PN). Due to lower sample sizes, LN subtypes were not tested against PN. Friedman’s test with Dunn’s correction was used for paired data. Error bars represent s.e.m. Calculation of ΔF/F, % ΔF/F, as well as statistical analysis was performed in GraphPad Prism 7, Ca^2+^ response amplitudes were calculated via Microsoft Excel, and R Studio version 1.3.1056 was used for data management.

## Data availability

Data used in this study are available on reasonable request.

## Acknowledgements

We like to thank Maja Bohn for help with the cell countings, and Michael Stern for his insightful comments to the manuscript. This research was partially funded by the German Federal Ministry of Education and Research (BMBF project 031L0062A).

## Author-Contributions

GAB conceived, performed, analyzed the experiments and wrote the paper with the help of GB. GB conceived and supervised the research.

## List of abbreviations

AL: Antennal lobe
ORN: Olfactory recetor neuron
PN: Projection neurons
LN: Local neurons
GABA-IR: GABA immunoreactivity
NADPHd: NADPH diaphorase
nAChR: Nicotinic acetylcholine receptor
mAChR: Muscarinic acetylcholine receptor
ROC: Reciever operating characteristic

## References

1 Wang, X. et al. The locust genome provides insight into swarm formation and long-distance flight. Nat Commun 5, 2957, doi:10.1038/ncomms3957 (2014).

2 Zhang, L., Lecoq, M., Latchininsky, A. & Hunter, D. Locust and Grasshopper Management. Annual Rev Entomol 64, 15–34 (2019).

3 Verlinden, H. et al. First draft genome assembly of the desert locust, Schistocerca gregaria. F1000Research 9, 775, doi:10.12688/f1000research.25148.1 (2020).

4 Guo, X. et al. 4-Vinylanisole is an aggregation pheromone in locusts. Nature 584, 584–588 (2020).

5 Burrows, M. Neurobiology of an Insect Brain. (Oxford University Press, 1996).

6 Roeder, T. Biochemistry and molecular biology of receptors for biogenic amines in locusts. Microsc Res Tech 56, 237–247 (2002).

7 Goodman, C. S. & Bate, M. Neuronal development in the grasshopper. Trends Neurosci 4, 163 – 169 (1981).

8 Truman, J., Ewer, J. & Ball, E. Dynamics of cyclic GMP levels in identified neurones during ecdysis behaviour in the locust Locusta migratoria. J Exp Biol 199, 749–758 (1996).

9 Laurent, G. Dynamical representation of odors by oscillating and evolving neural assemblies. Trends Neurosci 19, 489–496 (1996).

10 Hansson, B. S. & Stensmyr, M. C. Evolution of insect olfaction. Neuron 72, 698–711 (2011).

11 Vosshall, L. B., Amrein, H., Morozov, P. S., Rzhetsky, A. & Axel, R. A spatial map of olfactory receptor expression in the Drosophila antenna. Cell 96, 725–736 (1999).

12 Buck, L. & Axel, R. A novel multigene family may encode odorant receptors: a molecular basis for odor recognition. Cell 65, 175–187 (1991).

13 Anton, S. & Hansson, B. S. Antennal lobe interneurons in the desert locust Schistocerca gregaria (Forskal): processing of aggregation pheromones in adult males and females. J Comp Neurol 370, 85–96 (1996).

14 Ernst, K. D., Boeckh, J. & Boeckh, V. A neuroanatomical study on the organization of the central antennal pathways in insects. Cell Tissue Res 176, 285–306 (1977).

15 Seidel, C. & Bicker, G. Colocalization of NADPH-diaphorase and GABA-immunoreactivity in the olfactory and visual system of the locust. Brain Res 769, 273–280 (1997).

16 Leitch, B. & Laurent, G. GABAergic synapses in the antennal lobe and mushroom body of the locust olfactory system. J Comp Neurol 372, 487–514 (1996).

17 Laurent, G. Olfactory network dynamics and the coding of multidimensional signals. Nat Rev Neurosci 3, 884–895 (2002).

18 Laurent, G. & Naraghi, M. Odorant-induced oscillations in the mushroom bodies of the locust. J Neurosci 14, 2993–3004 (1994).

19 Lutz, E. M. & Tyrer, N. M. Immunohistochemical localization of choline acetyltransferase in the central nervous system of the locust. Brain Res 407, 173–179 (1987).

20 Knipper, M., Strotmann, J., Mädler, U., Kahle, C. & Breer, H. Monoclonal antibodies against the high affinity choline transport system. Neurochem Int 14, 217–222 (1989).

21 Rind, F. C. & Leitinger, G. Immunocytochemical evidence that collision sensing neurons in the locust visual system contain acetylcholine. J Comp Neurol 423, 389–401 (2000).

22 Ehrhardt, E. & Boyan, G. Evidence for the cholinergic markers ChAT and vAChT in sensory cells of the developing antennal nervous system of the desert locust Schistocerca gregaria. Invert Neurosci 20, 19, doi:10.1007/s10158-020-00252-4 (2020).

23 Breer, H. & Sattelle, D. B. Molecular properties and functions of insect acetylcholine receptors. J Insect Physiol 33, 771 – 790 (1987).

24 Gundelfinger, E. D. How complex is the nicotinic receptor system of insects? Trends Neurosci 15, 206–211 (1992).

25 Caulfield, M. P. Muscarinic receptors--characterization, coupling and function. Pharmacol Ther 58, 319–379 (1993).

26 Trimmer, B. A. Characterization of a muscarinic current that regulates excitability of an identified insect motoneuron. J Neurophysiol 72, 1862–1873 (1994).

27 Fiala, A. et al. Genetically expressed cameleon in Drosophila melanogaster is used to visualize olfactory information in projection neurons. Curr Biol 12, 1877–1884 (2002).

28 Wang, J. W., Wong, A. M., Flores, J., Vosshall, L. B. & Axel, R. Two-photon calcium imaging reveals an odor-evoked map of activity in the fly brain. Cell 112, 271–282 (2003).

29 Rozenfeld, E., Lerner, H. & Parnas, M. Muscarinic Modulation of Antennal Lobe GABAergic Local Neurons Shapes Odor Coding and Behavior. Cell Rep 29, 3253–3265.e4 (2019).

30 Moreaux, L. & Laurent, G. Estimating firing rates from calcium signals in locust projection neurons in vivo. Front Neural Circuits 1, 2, doi:10.3389/neuro.04.002.2007 (2007).

31 Gocht, D., Wagner, S. & Heinrich, R. Recognition, presence, and survival of locust central nervous glia in situ and in vitro. Microsc Res Tech 72, 385–397 (2009).

32 Trimmer, B. A. Current excitement from insect muscarinic receptors. Trends Neurosci 18, 104–111 (1995).

33 Bicker, G. & Kreissl, S. Calcium imaging reveals nicotinic acetylcholine receptors on cultured mushroom body neurons. J Neurophysiol 71, 808–810 (1994).

34 Cayre, M., Buckingham, S. D., Yagodin, S. & Sattelle, D. B. Cultured insect mushroom body neurons express functional receptors for acetylcholine, GABA, glutamate, octopamine, and dopamine. J Neurophysiol 81, 1–14 (1999).

35 MacLeod, K. & Laurent, G. Distinct mechanisms for synchronization and temporal patterning of odor-encoding neural assemblies. Science 274, 976–979 (1996).

36 Hermsen, B. et al. Neuronal nicotinic receptors in the locust Locusta migratoria. Cloning and expression. J Biol Chem 273, 18394–18404 (1998).

37 Marshall, J. et al. Sequence and functional expression of a single alpha subunit of an insect nicotinic acetylcholine receptor. Embo j 9, 4391–4398 (1990).

38 Amar, M., Thomas, P., Wonnacott, S. & Lunt, G. G. A nicotinic acetylcholine receptor subunit from insect brain forms a non-desensitising homo-oligomeric nicotinic acetylcholine receptor when expressed in Xenopus oocytes. Neurosci Lett 199, 107–110 (1995).

39 Jones, A. K. et al. Sgbeta1, a novel locust (Schistocerca gregaria) non-alpha nicotinic acetylcholine receptor-like subunit with homology to the Drosophila melanogaster Dbeta1 subunit. Invert Neurosci 5, 147–155 (2005).

40 Breer, H. & Knipper, M. Characterization of acetylcholine release from insect synaptosomes. Insect Biochemistry 14, 337 – 344 (1984).

41 Knipper, M. & Breer, H. Subtypes of muscarinic receptors in insect nervous system. Comp Biochem Physiol C 90, 275 – 280 (1988).

42 Leitch, B. & Pitman, R. M. Modulation of transmitter release from the terminals of the locust wing stretch receptor neuron by muscarinic antagonists. J Neurobiol 28, 455–464 (1995).

43 Judge, S. & Leitch, B. Modulation of transmitter release from the locust forewing stretch receptor neuron by GABAergic interneurons activated via muscarinic receptors. J Neurobiol 40, 420–431 (1999).

44 Oertner, T. G., Single, S. & Borst, A. Separation of voltage- and ligand-gated calcium influx in locust neurons by optical imaging. Neurosci Lett 274, 95–98 (1999).

45 Campusano, J. M., Su, H., Jiang, S. A., Sicaeros, B. & O’Dowd, D. K. nAChR-mediated calcium responses and plasticity in Drosophila Kenyon cells. Dev Neurobiol 67, 1520–1532 (2007).

46 Müller, U. & Bicker, G. Calcium-activated release of nitric oxide and cellular distribution of nitric oxide-synthesizing neurons in the nervous system of the locust. J Neurosci 14, 7521–7528 (1994).

47 Bicker, G., Schmachtenberg, O. & Vente, J. D. Geometric considerations of nitric oxide-cyclic GMP signalling in the glomerular neuropil of the locust antennal lobe. Proc R Soc Lond B 264, 1177–1181 (1997).

48 Jiang, S. A., Campusano, J. M., Su, H. & O’Dowd, D. K. Drosophila mushroom body Kenyon cells generate spontaneous calcium transients mediated by PLTX-sensitive calcium channels. J Neurophysiol 94, 491–500 (2005).

49 Bentley, D., Guthrie, P. B. & Kater, S. B. Calcium ion distribution in nascent pioneer axons and coupled preaxonogenesis neurons in situ. J Neurosci 11, 1300–1308 (1991).

50 Isaacson, M. D. & Hedwig, B. Electrophoresis of polar fluorescent tracers through the nerve sheath labels neuronal populations for anatomical and functional imaging. Scientific Reports 7, 40433, doi:10.1038/srep40433 (2017).

51 Li, Y. et al. CRISPR/Cas9 in locusts: Successful establishment of an olfactory deficiency line by targeting the mutagenesis of an odorant receptor co-receptor (Orco). Insect Biochem Mol Biol 79, 27–35 (2016).

52 Bicker, G. & Stern, M. Structural and Functional Plasticity in the Regenerating Olfactory System of the Migratory Locust. Front Physiol 11, 608661, doi:10.3389/fphys.2020.608661 (2020).

53 Bergmann, G. A. et al. An intact insect embryo for developmental neurotoxicity testing of directed axonal elongation. ALTEX 36, 643–649 (2019).

54 Bode, K. et al. A locust embryo as predictive developmental neurotoxicity testing system for pioneer axon pathway formation. Arch Toxicol 94, 4099–4113 (2020).

55 Kirchhof, B. & Bicker, G. Growth properties of larval and adult locust neurons in primary cell culture. J Comp Neurol 323, 411–422 (1992).

56 Bicker, G., Naujock, M. & Haase, A. Cellular expression patterns of acetylcholinesterase activity during grasshopper development. Cell Tissue Res 317, 207–220 (2004).

57 Stern, M. et al. Development of nitrergic neurons in the nervous system of the locust embryo. J Comp Neurol 518, 1157–1175 (2010).

58 Abercrombie, M. Estimation of nuclear population from microtome sections. Anat Rec 94, 239–247 (1946).

59 Schindelin, J. et al. Fiji: an open-source platform for biological-image analysis. Nat Methods 9, 676–682 (2012).

