## Supplemental Material for "Cholinergic calcium responses in cultured antennal lobe neurons of the migratory locust"

**Supplementary Table S1: Collection of calcium indicators used for screening.** Calcium indicators were dissolved in DMSO and diluted to appropriate concentrations in Leibovitz's L15 medium (maximum DMSO concentration 0.1 %). Incubation periods were 30 min, 1 h, 3 h and 24 h for each indicator followed by a washout. After addition of 5 µM calcimycine, a pronounced increase in fluorescence was only achieved using 1 h incubation in 1 µM Cal 520-AM. All other listed calcium indicators failed in this explorative trial.

| Calcium indicator | Test Conc. | Loading | ΔF increase | Supplier |
|---|---|---|---|---|
| Cal 520-AM | 1, 5, 8, 10, 15 µM | 30 min – 24 h | + | Abcam ab171868 |
| Calcium Green-1 Dextran 3000 MW | 1, 5 µM | 30 min – 24 h | - | Molecular Probes C6765 |
| Calcium Green-1-AM | 1, 5 µM | 30 min – 24 h | - | Molecular Probes C3012 |
| Calcium Green-2-AM | 1, 5 µM | 30 min – 24 h | - | Molecular Probes C3732 |
| Calcium Orange-AM | 1, 5 µM | 30 min – 24 h | - | Molecular Probes C3732 |
| Fluro-3-AM | 1, 5 µM | 30 min – 24 h | - | Invitrogen F1242 |

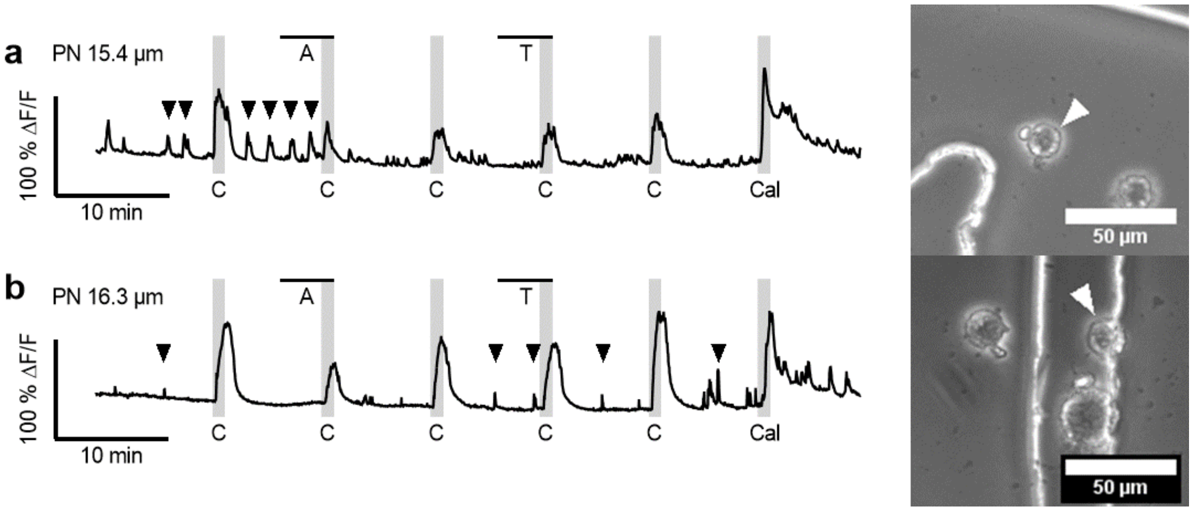


**Supplementary Figure S2: Spontaneous calcium transients of isolated antennal lobe neurons.** Calcium signal traces of cultured projection neurons (white arrowhead) are shown (**a** & **b**). Cultured antennal lobe neurons display spontaneous calcium transients (black arrowheads), which were lower in magnitude and shorter in duration than transients elicited via pharmacological stimulation. We found no neuronal connections between these examples and adjacent neurons. Spontaneous transients were not consistent in magnitude, duration, and frequency between neurons, nor throughout the imaging period (**a** & **b**). In **a** repetitive spontaneous transients were observed prior to and after stimulation with carbachol. In **b**, spontaneous transients were lower in magnitude and irregular in frequency throughout the imaging period. Spontaneous transients were still discernable during preincubation with either cholinergic antagonist. Neither atropine nor tubocurarine had apparent effects on spontaneous transients during preincubation. Definitive effects of agonists or antagonists on spontaneous transient generation were not ascertained in this study and may be subject to further, more detailed investigations.

**Supplementary Video S3: Time lapse video of cultivated antennal lobe neurons, loaded with the calcium sensor Cal-520.** Cells of the $Ca^{2+}$ imaging traces depicted in Fig.5a i - iv (**a** - **d** in same order as in Fig.5a) are marked in white circles. Fluorescence intensity is shown in false color (LUT between 0 and 255). Application of stimulants (e.g. Nicotine) is represented in a color change of the respective label from white to orange for the duration of the stimulus. Surrounding cells also show occasional spontaneous activity before and in between stimulations.

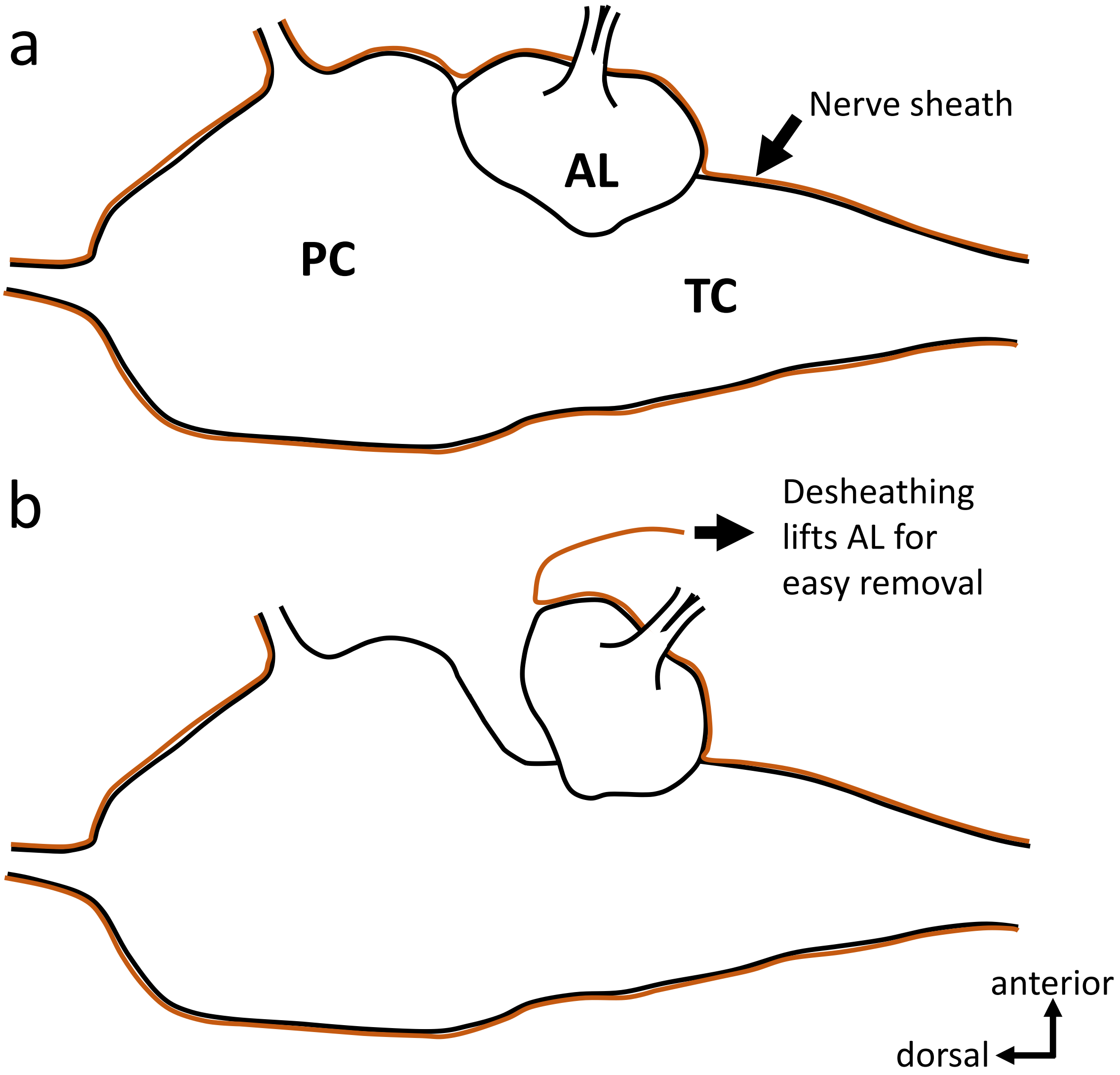


**Supplementary Figure S4: Schematic side view of the locust brain before (a) and during desheathing (b).** The antennal lobe (AL) is attached to the protocerebrum (PC) and the tritocerebrum (TC). There are no nerve tracts running towards the PC on its dorsally oriented side. When the nerve sheath is pulled from the PC towards the TC, this arrangement results in an uplift, allowing for easy removal of the AL and its cells.
